# Comprehensive Molecular Characterization of High-Grade Endometrial Cancer in an Ancestrally Diverse Cohort

**DOI:** 10.64898/2026.05.01.721962

**Authors:** Marina Frimer, Devin Gee, Zoe R. Goldstein, William F. Hooper, Kyriaki Founta, Astrid Deschênes, Heather Geiger, Pascal Belleau, Melissa Kramer, Brian Yueh, Tim Chu, Ali Oku, Zalman Vaksman, Valentina Grether, Zoe Steinsnyder, Andrew L. Araneo, Charlie Chung, Arisa Kapedani, Aaron Nizam, Onur Eskiocak, Kadir Ozler, Gary L. Goldberg, Alexander Krasnitz, W. Richard McCombie, Mali Barbi, Lara Winterkorn, Nicolas Robine, Semir Beyaz, Nyasha Chambwe

## Abstract

Endometrial cancer (EC) exhibits one of the most striking racial disparities in oncology with black women disproportionately affected by aggressive high-grade subtypes that have poorer outcomes. While social and environmental factors undoubtedly contribute, the molecular underpinnings of these disparities remain critically understudied. To bridge this knowledge gap, we performed matched tumor-normal whole-genome sequencing and tumor transcriptome sequencing on 71 predominantly high-grade EC patient samples from an ancestrally diverse cohort of women recruited at a large hospital system in the New York metropolitan area. Our analysis characterized the germline and somatic mutation landscape, identifying ancestry-associated molecular differences. Notably, focal amplification of the *EVI1* transcription factor (encoded at the MECOM locus) was significantly more frequent in African ancestry patients and associated with poorer clinical outcomes in an external validation cohort. Additionally transcriptome analysis revealed decreased CD8+ T cell infiltration with increasing African ancestry, suggesting tumor immune microenvironment differences with potential therapeutic implications.

**Graphical Abstract:** 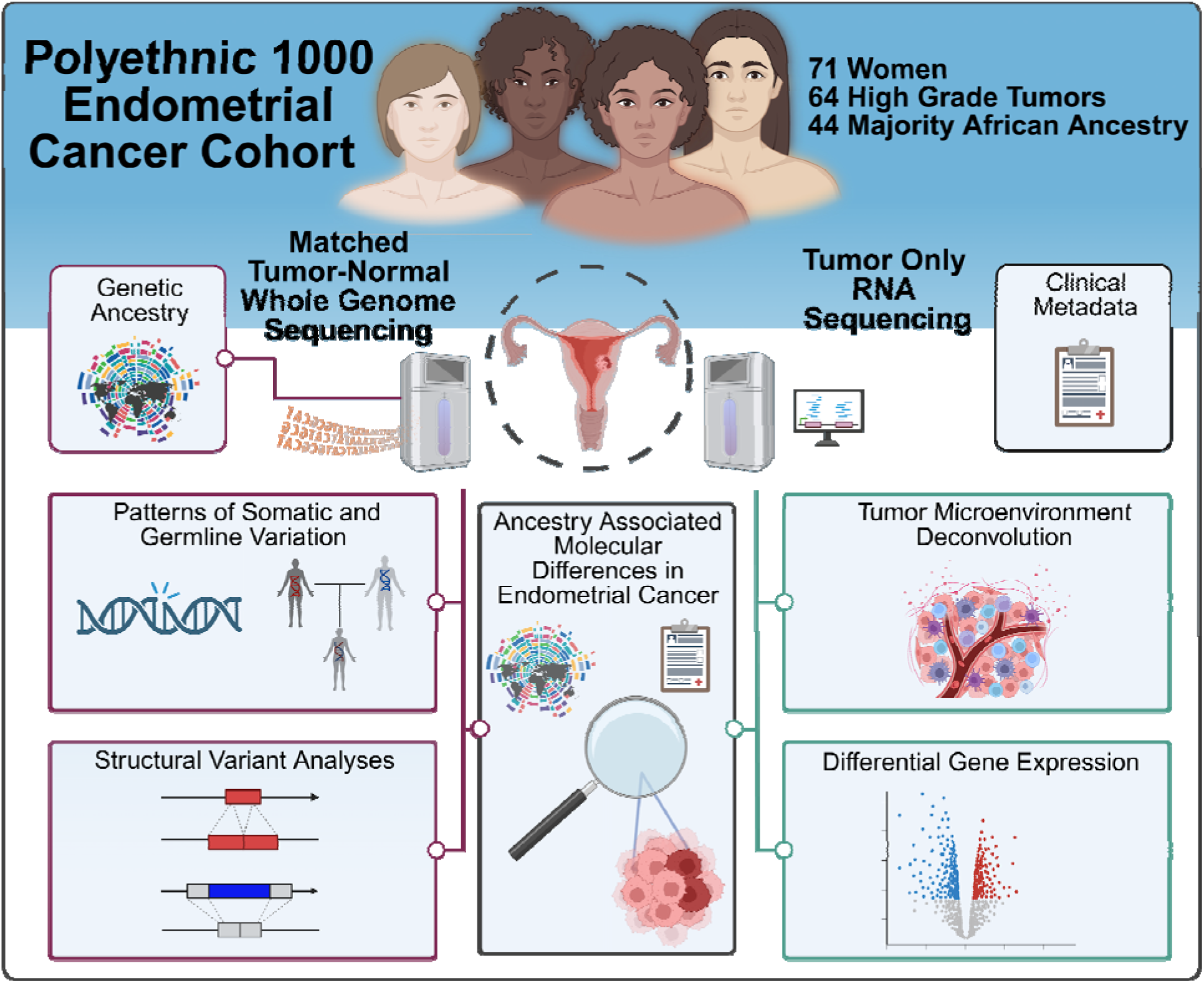

**Highlights:**

- This study represents the most ancestrally diverse whole-genome sequencing characterization of high-grade endometrial cancer, with 62% of patients of African ancestry.
- MECOM focal amplification preferentially targets the oncogenic short isoform (EVI1) and is more frequent in patients of African ancestry.
- African ancestry is associated with reduced CD8+ T cell infiltration and differential activation of immune and metabolic pathways in copy-number high endometrial tumors.

## Introduction

Endometrial cancer (EC) is the most common gynecologic malignancy diagnosed in the United States with an estimated 69,120 new cases and 13,860 deaths in 2025.^1^ Despite recent epidemiological trends indicating declining overall cancer mortality, both EC incidence and mortality are increasing, with projections of 120,000 new cases and 24,000 deaths per year by 2050.^2^ Of critical concern are the pronounced racial and ethnic disparities in EC burden with larger increases in incidence and mortality observed in self-identified black or African American (B/AA) women compared to white women.^3^ Alarmingly, EC incidence is projected to increase disproportionately in B/AA women by 2050 with rates in white women projected to increase from 57.7 to 74.2 cases per 100,000, while in B/AA women the projected increase is from 56.8 to 86.9 per 100,000.^2^

Although most ECs (84%) are the less aggressive low-grade type I cancers with endometrioid histology,^4^ B/AA women are more likely to be diagnosed with type II cancers that are predominantly of serous, clear-cell or carcinosarcoma histology.^5^ These type II cancers represent ∼10% of all EC, but account for ∼40% of all EC deaths.^6^ These aggressive histologic subtypes are significantly more likely to present at later stages with extra-uterine disease, exhibit poor responses to chemotherapy or radiation, carry a worse prognosis.^4,7^ The higher EC-related disease burden in B/AA women can be attributed, at least in part, to this population being less likely to receive guideline-compliant treatment,^3^ namely, B/AA women are less likely to receive surgery, to have minimally invasive surgery, less likely to have lymph node sampling/dissection, or to receive chemotherapy.^8^ Clinical trial participation also reveals stark disparities, with African American women comprising only 5% of patients in phase I gynecologic oncology trials. Participation among African American women decreased from 11.4% in 1995-1999 to 6.2% in 2015-2018, representing a 1.8-fold decline.^8^

While The Cancer Genome Atlas (TCGA) has provided transformative insights into EC molecular heterogeneity by identifying the four molecular subtypes: POLE ultramutated, microsatellite instability hypermutated, copy-number low (CN-L), and copy-number high (CN-H) tumors — its representation of B/AA women remains critically low, with approximately 12% of the cohort identifying as B/AA compared to 78% white. This lack of diversity limits the applicability of TCGA findings to understanding disparities in B/AA women, whose tumors may exhibit unique molecular characteristics.^9^

Despite these known challenges, there has been limited progress in addressing the disproportionate disease burden faced by B/AA women. There is an urgent need for effective therapeutic development and comprehensive research initiatives to resolve this inequity. Our study aims to fill these gaps by focusing on the genetic and molecular factors associated with aggressive subtypes of EC and the observed disparities. To achieve these goals, we established a robust clinical and experimental framework to investigate the genetic and molecular basis of EC disparities in partnership with the Polyethnic-1000 initiative (P-1000). P-1000 is a collaborative effort organized by the New York Genome Center and leading local cancer research institutions to advance cancer genomics in diverse participants.^10^ Our efforts included targeted recruitment of participants from historically underrepresented racial and ethnic groups and the creation of a well-annotated, racially diverse EC biobank with biospecimens from those individuals.

Here we describe the whole genome and transcriptome characterization of 71 women diagnosed with high-grade EC histological subtypes. To address the critical limitations of previous studies, which have relied predominantly on exome or panel-based sequencing in cohorts with limited ancestral diversity, our study employs matched tumor-normal whole genome sequencing (WGS) in a cohort enriched for women self-identifying as B/AA. This approach enables the characterization of the complete landscape of somatic alterations, including single nucleotide variants, structural variants, and both large-scale and focal copy number events. Our analysis reveals that focal amplification of the oncogenic EVI1 isoform at the MECOM locus is significantly more frequent in patients of African ancestry and is associated with worse clinical outcomes, a finding independently validated in reprocessed TCGA WGS data. Complementary transcriptomic analyses revealed ancestry-associated differences in pathway activation and decreased CD8+ T cell infiltration with increasing African ancestry, suggesting that both tumor-intrinsic and microenvironmental features may contribute to the disparities observed in high-grade EC.

## Results

### Study overview

To better understand genetic and molecular drivers of high-grade EC in the context of racial and ethnic disparities, we performed matched tumor-normal whole genome sequencing (WGS) and tumor-only bulk transcriptome sequencing (RNA-seq) on EC samples from a racially diverse cohort of women (**Fig. 1a**). We sequenced 78 tumor and normal samples to a median coverage of 94X and 47X, respectively and confirmed the absence of significant tumor-in-normal and inter-individual contamination that would hamper downstream analysis resulting in 71 eligible samples (**Supplementary Fig. 1 and 2; Supplementary Table S1)**. 90% of the cohort was made up of grade III tumors although the tumor stage at diagnosis varied (**Table 1**). The median age at diagnosis was 68 and median body mass index (BMI) was 31. 75% (53) of the women in the study self-declared as B/AA. Estimated global genetic ancestry from germline matched samples (see **Methods**) revealed that the cohort included 44 (62%) individuals with predominant African ancestry (AFR) (admixture coefficient ≥80%) **(Fig. 1b-c; Table 1; Supplementary Table S2)**, in line with 73% who self-identified as B/AA by race. 9 individuals had majority European ancestry (EUR), 4 with majority South Asian (SAS), 2 East Asian (EAS), 1 Admixed American (AMR) and 7 mixed ancestry (Admixed, no dominant continental ancestry ≥80%) **(Fig. 1c; Table 1)**. In contrast to prior studies, this EC cohort is significantly enriched for women of West African ancestry and offers the potential to characterize ancestry-associated molecular disease drivers in the more aggressive ECs represented in this cohort **(Fig. 1b; Supplementary Fig. 3; Supplementary Table S2)**.

**Fig. 1:**
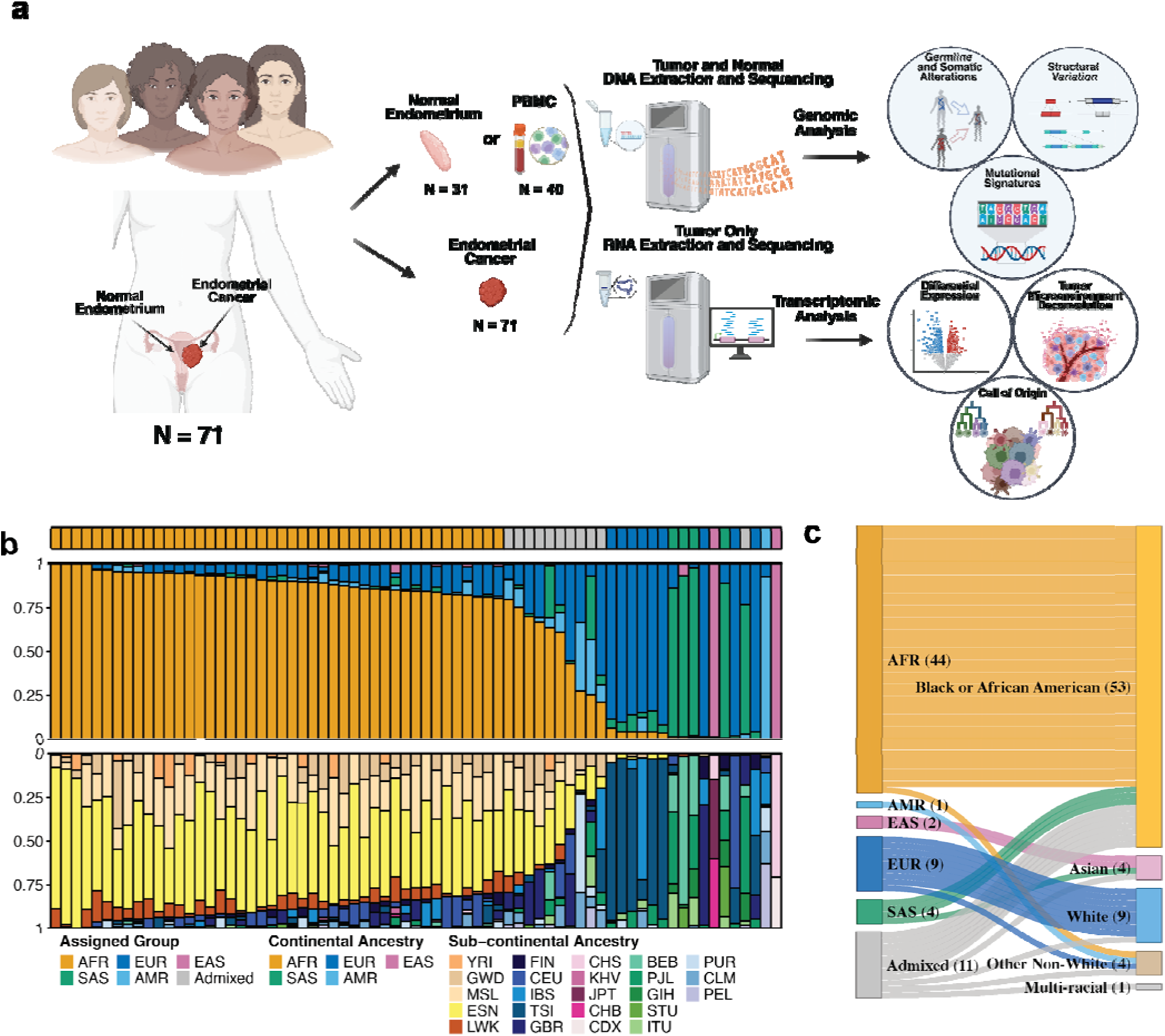
Study design and summary of estimated global genetic ancestry of the study cohort. **a,** Experimental workflow schematic showing an overview of biospecimen acquisition, DNA and RNA extraction, sequencing and bioinformatic analysis. **b,** Distribution of the patients per assigned continental ancestry (n=71). Inferred continental and sub-continental admixtures in the cohort with the assigned continental ancestry in the top track. **c,** The relationship between assigned continental genetic ancestry (on the left) and self-identified race (on the right) for the cohort.

**Fig. 2:**
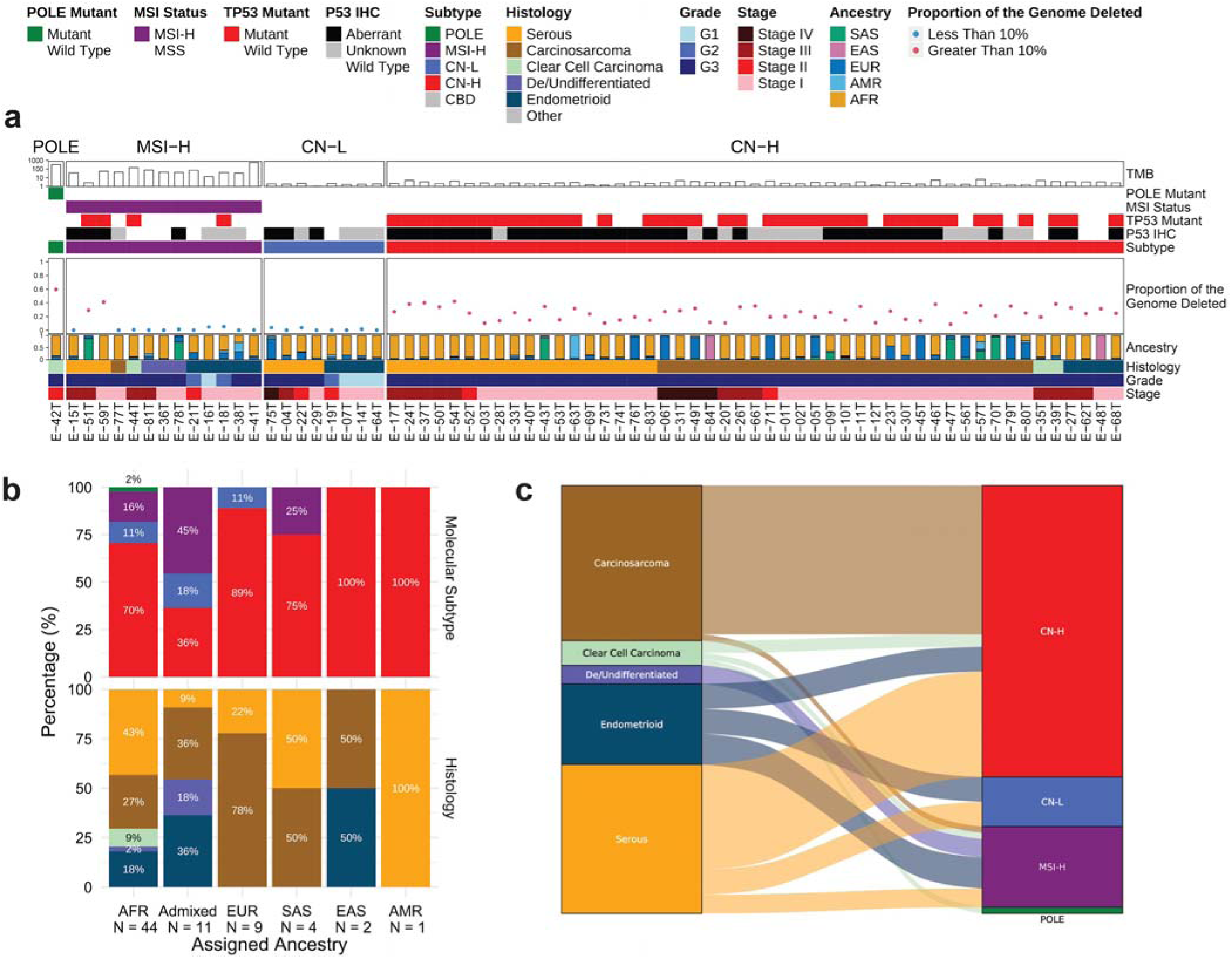
Overview of molecular and histological subtypes represented in the study cohort. **a**, Samples were classified as POLE mutant (POLE), Microsatellite Instability High (MSI-H), Copy Number Low (CN-L), and Copy Number High (CN-H) based on somatic mutations in the POLE exonuclease domain, p53 and mismatch repair pathway immunohistochemistry, computationally derived microsatellite instability, and copy number variability analyses. The molecular subtypes and the associated characteristics are summarized over histology, grade, and stage in the annotations below. **b,** The relative distribution of molecular(top) and histological (bottom) subtypes across the cohort. **c,** Sankey plot displaying molecular subtype and histology relationships across the cohort

**Table 1:**
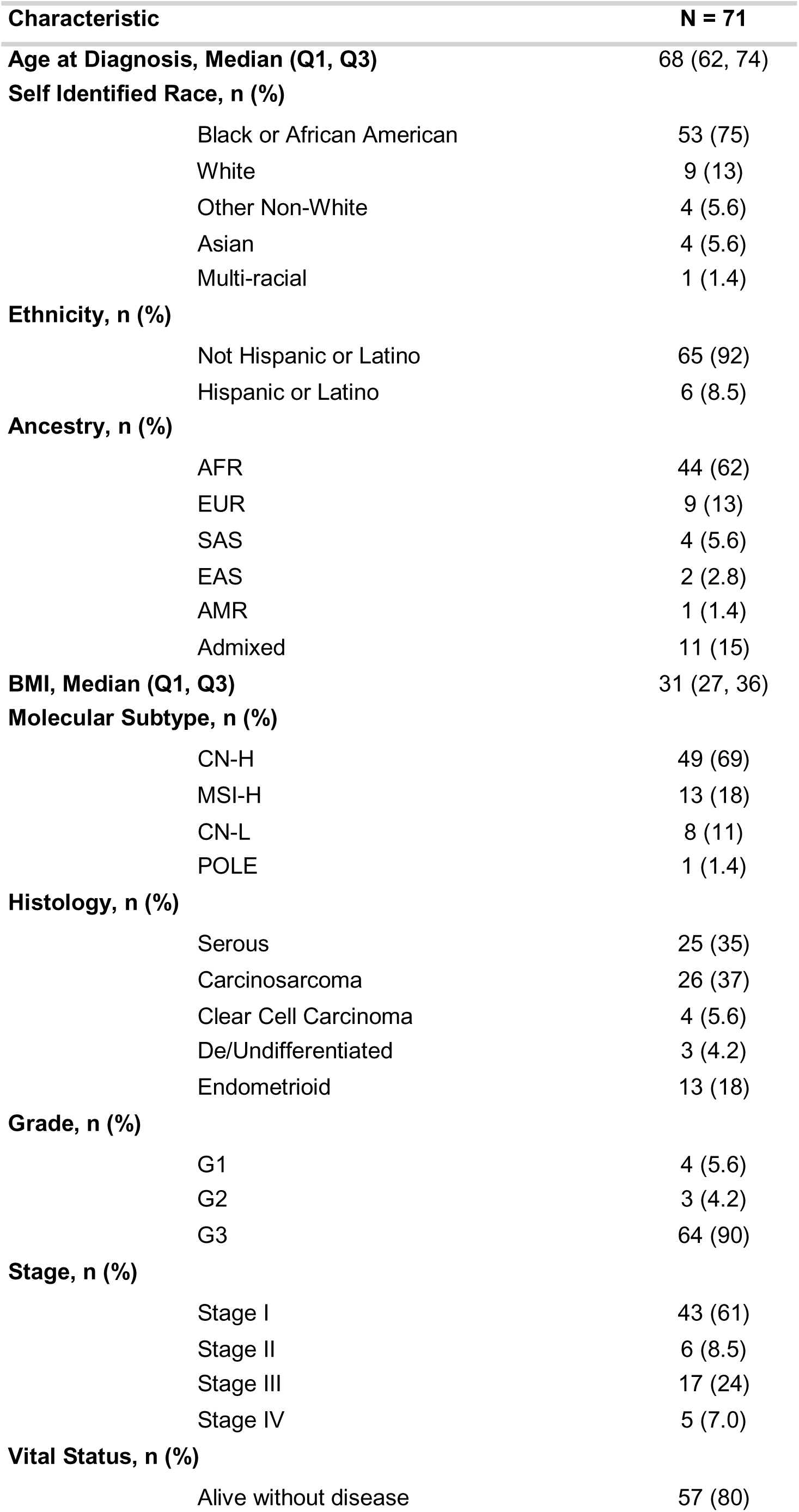
Overview of Clinical and Demographic Characteristics of the Cohort.

| Characteristic | N = 71 |
| --- | --- |
| **Age at Diagnosis, Median (Q1, Q3)** | 68 (62, 74) |
| **Self Identified Race, n (%)** |  |
| Black or African American | 53 (75) |
| White | 9 (13) |
| Other Non-White | 4 (5.6) |
| Asian | 4 (5.6) |
| Multi-racial | 1 (1.4) |
| **Ethnicity, n (%)** |  |
| Not Hispanic or Latino | 65 (92) |
| Hispanic or Latino | 6 (8.5) |
| **Ancestry, n (%)** |  |
| AFR | 44 (62) |
| EUR | 9 (13) |
| SAS | 4 (5.6) |
| EAS | 2 (2.8) |
| AMR | 1 (1.4) |
| Admixed | 11 (15) |
| **BMI, Median (Q1, Q3)** | 31 (27, 36) |
| **Molecular Subtype, n (%)** |  |
| CN-H | 49 (69) |
| MSI-H | 13 (18) |
| CN-L | 8 (11) |
| POLE | 1 (1.4) |
| **Histology, n (%)** |  |
| Serous | 25 (35) |
| Carcinosarcoma | 26 (37) |
| Clear Cell Carcinoma | 4 (5.6) |
| De/Undifferentiated | 3 (4.2) |
| Endometrioid | 13 (18) |
| **Grade, n (%)** |  |
| G1 | 4 (5.6) |
| G2 | 3 (4.2) |
| G3 | 64 (90) |
| **Stage, n (%)** |  |
| Stage I | 43 (61) |
| Stage II | 6 (8.5) |
| Stage III | 17 (24) |
| Stage IV | 5 (7.0) |
| **Vital Status, n (%)** |  |
| Alive without disease | 57 (80) |

The study cohort’s heterogeneity extends further to the histological and molecular subtypes. As summarized in (**Fig. 2a)**, we categorized the 71 primary tumor samples into the four molecular subtypes first defined in the TCGA endometrial carcinoma cohort (UCEC)^9^ by leveraging our WGS dataset to determine POLE mutation and microsatellite instability (MSI) status. We performed orthogonal validation of the WGS-derived MSI status call with clinical immunohistochemistry (IHC) testing data where available **(Supplementary Fig. 4)**. 69% of the tumors were CN-H, 11.3% were CN-L, 18.3% were MSI-H, and 1.4% were POLE **(Fig. 2a; Table 1)**. While we did not observe a significant association between molecular subtype and genetic ancestry (p = 0.467, Fisher’s exact test) (**Supplementary Table S3**), the majority of women of AFR ancestry in our cohort had CN-H tumors (70.45%), a finding that has been previously reported.^11,12^ Each assigned genetic ancestry group was represented among individuals with CN-H tumors powering the investigation of ancestry associated differences in the most aggressive subtype.

From a histological perspective, 82% of the tumors in the cohort were of non-endometrioid subtypes, primarily serous (35%) and carcinosarcoma (37%), with a small number of clear cell carcinoma and de/undifferentiated. There was no significant association observed between histology and genetic ancestry in the cohort (p = 0.095, Fisher’s exact test) (**Fig. 2b; Supplementary Table S3**). Although the majority of carcinosarcoma and serous tumors are of the CN-H subtype, there are complex interactions between histology and molecular subtype (**Fig. 2c**). This cohort stands in stark contrast to other genomically characterized EC cohorts such as those from TCGA,^9,13^ CPTAC^14–16^ or clinically ascertained cohorts like those from Weigelt et al.,^11^ in that it is strongly enriched in CN-H tumors with primarily serous/carcinosarcoma histology sourced from a majority of B/AA women (**Supplementary Fig. 5).** In summary, the demographic, clinical and molecular characteristics of this cohort present the opportunity to discover novel oncogenic drivers of the more aggressive subtypes of EC.

### Germline predisposition variant analysis

We next identified disease-associated pathogenic or likely pathogenic (P/LP) germline variants of interest, limiting our analysis to 465 genes annotated as cancer predisposition genes^17,18^ or involved in DNA repair^19^ (**Supplementary Table S4**) due to the relatively small size of our cohort. 38% of the cohort (27/71 individuals) harbored one or more germline P/LP variant(s), of which 20/27 (74.1%) had at least one P/LP variant represented in ClinVar^20^ (**Supplementary Table S5**). Stop gain pathogenic variants were identified in mismatch repair genes linked to Lynch Syndrome, the most common hereditary syndrome associated with EC,^21–23^ with 2 *MSH2* and 1 *MSH6* pathogenic variant carriers in the cohort. In addition, we detected P/LP variants in *BRCA1*, *PALB2*, and *BRIP1;* genes involved in DNA double strand break repair and implicated in hereditary breast and ovarian cancer syndrome.^20,24^ Other potential EC predisposition variants were found in the DNA repair genes *LIG4*, *PNKP*, *ATR* and other cancer predisposition genes *ELANE* and *MYLK*. Our findings revealed that although P/LP germline variants conferring disease predisposition were present in a subset of individuals. No particular genes with such variants were enriched in any ancestry or molecular subgroup within this cohort although the sample size in this cohort makes it infeasible to detect enrichment.

### The landscape of ancestry-associated somatic genetic alterations

We observed a median somatic tumor mutation burden (TMB) of 3.1 somatic mutations per megabase (mut/Mb, range 1.2-634.6). As expected, TMB was significantly elevated in MSI-H tumors (median 49.6 mut/Mb) as compared to CN-L and CN-H (p-value<0.001, Wilcoxon rank sum test). When controlling for molecular subtype, TMB was not significantly different between ancestries or histologies (**Supplementary Fig. 6**).

The most frequently altered genes in our cohort were known EC drivers such as *TP53* (42/71, 59%), *PIK3CA* (25/71, 35%), *ARID1A* (20/71, 28%), and *PTEN* (18/71, 25%)(**Fig. 3a**), similar to TCGA EC cohorts^9,13^. As expected, *TP53* was most frequently mutated in carcinosarcoma and serous tumors, while *PIK3CA*, *PTEN*, and *ARID1A* mutational frequencies were higher in endometrioid, clear cell carcinoma, and dedifferentiated adenocarcinoma (**Fig. 3a**). Stratifying by molecular subtype, *TP53* was most frequently mutated in CN-H (38/49, 78%), accompanied by loss of heterozygosity in all but one case. Among the 11 CN-H tumors with no *TP53* mutations, 5 had an *ATM* mutation, confirming a previously reported pattern of mutual exclusivity^25^. *ARID1A* and *PIK3CA* were mutated in 3/8 CN-L samples, and the remaining 5 samples had no nonsynonymous mutations in *TP53*, *ARID1A*, *PIK3CA*, and *PTEN*. *SETD1B* (12/13, 92%) and *KMT2D* (10/13, 77%) were highly mutated in MSI-H tumors, in addition to the aforementioned *ARID1A* (10/13, 77%), *PIK3CA* (8/13, 62%), and PTEN (8/13, 62%).

**Fig. 3:**
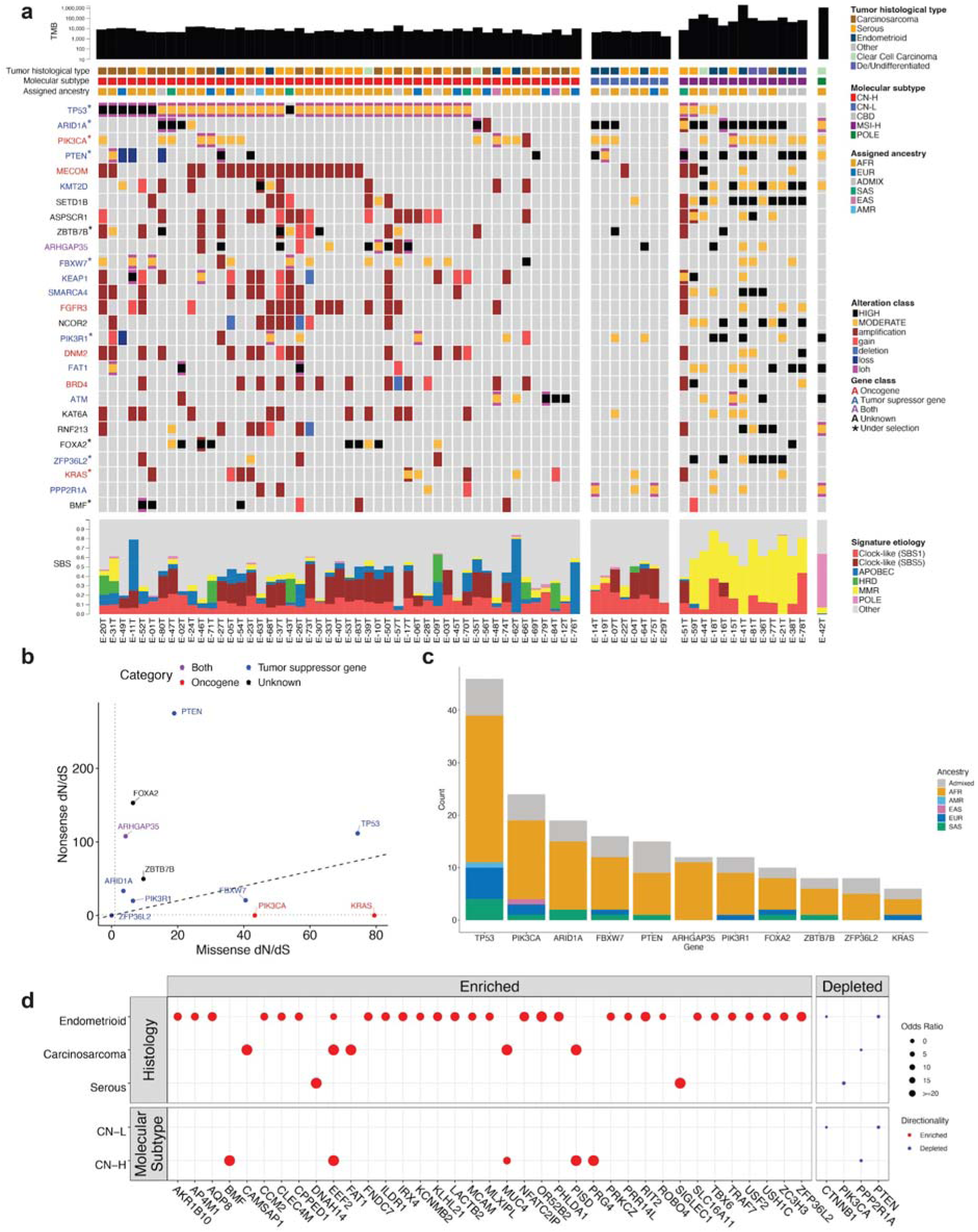
Somatic mutation landscape of predominantly high-grade endometrial cancer. **a,** Summary of alterations and mutational signatures across samples in the cohort, grouped by molecular subtype. The top row corresponds to the number of somatic SNV/indels, represented in log_10_ scale. Tumor histology, molecular subtype, and patients’ genetic ancestry are indicated below. In the middle, an oncoprint summarizes the mutational pattern of the most frequently mutated COSMIC Cancer Gene Census genes and genes under significant positive selection. The bottom panel shows the proportion of the most representative COSMIC Single Base Substitution signatures grouped by etiology. **b,** Maximum likelihood estimates of dN/dS ratios genes under significant positive selection. Point and text color indicates driver gene classification using OncoKB. **c,** Ancestry distributions of mutations in genes under significant positive selection. **d,** AFR group comparison by histology and molecular subtype to TCGA EUR and MSKCC white cohorts. Genes more frequently mutated in AFR cohort samples are “enriched” (p<0.05 unadjusted) and denoted by red dots. Genes less frequently mutated in the AFR cohort are “depleted” (p<0.05 unadjusted) and denoted by a blue dot. The size of each dot corresponds to the odds ratio.

We also noted an absence of *PPP2R1A* mutations in carcinosarcoma, in contrast to the 28% reported in TCGA-UCS, but in agreement with an earlier study^26^. The *PPP2R1A* mutation rate in other histologies (15.6%) was broadly in line with TCGA-UCEC (9.5%) and previous reports from MSKCC (16.3%)^27^ and CPTAC (9.5%).^16^ The ubiquitin ligase *FBXW7* was amplified or mutated in 23% (10/44) patients of African ancestry, *FBXW7* was previously detected in a large pancancer study of genetic ancestry as the single gene more frequently mutated in patients of African ancestry.^28^ In a recent meta-analysis of over 275,000 tumors, this enrichment was consistently significant in both endometrial and colorectal cancers.^29^

To differentiate between driver and passenger mutation events, we looked for signals of genes under positive selection as estimated by the ratio of nonsynonymous to synonymous mutations (**Methods**). We detected 11 genes under significant positive selection (FDR-adjusted p-value < 0.05) **(Fig. 3b; Supplementary Table S6)**. Of these genes, 9 were reported in a re-analysis of TCGA-UCEC with the same algorithm;^30^ *ZBTB7B* and *ZFP36L2* were novel, however, *ZBTB7B* was reported as being mutated in 8.77% of TCGA-UCS and 5.67% of TCGA-UCEC (respectively 15.3% in carcinosarcomas and 8.8% non-carcinosarcomas in our cohort). *ZFP36L2* was predominantly mutated in MSI-H tumors, suggesting that its significance could be driven by the overall high mutation burden in these samples. Adjusting for molecular subtype, no genes were found to be statistically more mutated in patients of African Ancestry compared to patients of all other ancestries (Cochran-Mantel-Haenszel test) (**Fig. 3c**).

As expected from Martincorena et al.,^30^ oncogenes tended to have a higher rate of (presumably activating) missense mutations, and tumor suppressors had a higher rate of truncating mutations. The tumor suppressor *FBXW7* was the exception, in that we detected a high rate of missense mutations in this gene. This phenomenon has been previously reported, wherein *FXBW7* missense mutations in WD40 domains essential for substrate recognition have a dominant negative effect on *FBXW7* homodimer function.^31,32^ Indeed, missense *FBXW7* mutations observed in our cohort tended to cluster in WD40 domain arginine residues (**Supplementary Fig. 7**). Of note, some of the genes detected as recurrently mutated in this cohort, such as *PIK3CA*, *PIK3R1*, *ARHGAP35*, *FBXW7*, *FOXA2*, *SPOP* have been reported as frequently mutated in normal endometrial glands.^33^

We observed significant ancestry-associated differences in the EC somatic alteration frequency when comparing AFR ancestry tumors in our cohort, with EUR ancestry tumors from the TCGA and the Weigelt et al. cohort ^11^. Tumors from AFR patients demonstrated a higher frequency of mutations in key genes, including *FAT1* and *EEF2* in carcinosarcomas and *ARID2* and *ARHGAP35* in endometrioid subtypes, while the CN-H subtype was enriched for alterations in *BMF*, *PISD*, and *PRG4*. Conversely, AFR tumors showed a significantly lower frequency of mutations in critical genes like *PTEN*, *CTNNB1*, and *PIK3CA* (**Fig. 3d, Supplementary Fig. 8**). Using annotations from MSKCC’s OncoKB^34,35^ and combining TCGA UCEC and our cohort (excluding carcinosarcomas), we observed a significant enrichment in clinically actionable mutations in patients of European ancestry compared to patients of African ancestry (p-value<0.001, Wilcoxon rank sum test) (**Supplementary Fig. 9**).

We decomposed the mutational patterns into the reference Catalogue of Somatic Mutations in Cancer (COSMIC)^36^ mutational signatures **(Fig. 3a; Supplementary Fig. 10; Supplementary Table S7)**. Almost all samples in the MSI-H subtype presented with SBS signatures associated with mismatch repair (MMR) deficiencies (SBS6, SBS14, SBS15, SBS20, SBS21, SBS26, SBS44) and a high proportion of the MMR-associated ID2 signature, as expected. We found that SBS1 was almost ubiquitous in the cohort, and that several samples presented with the other “clock-like signature” (SBS5), as well as some with signatures associated with APOBEC (SBS2, SBS13), and homologous recombination deficiency (SBS3), including in one serous and six carcinosarcoma cases. APOBEC-and HRD-associated signatures were more prevalent in the CN-H molecular subtype, present in 8.2% (4/49) and 14% (7/49) of the subtype respectively, and were not substantially present in the rest of the cohort **(Fig. 3a)**. Of the CN-H cases with substantial SBS3 detection, only two had concomitant evidence of microhomology mediated end-joining (ID6). ID6 has been shown to have more predictive value than SBS3 when nominating samples as homologous repair deficient via mutational signatures.^37^

Collectively these data suggest broad similarities in EC biology regardless of ancestry, with some interesting and novel findings in our cohort. Larger studies would be needed to fully characterize the effect of germline variation on the somatic profiles of EC tumors.

### Somatic copy number alterations in diverse ancestries

Our WGS-based approach enabled deep characterization of copy number and structural variation in this cohort. We applied the JaBbA algorithm^38^ to integrate read depth and breakpoints (hereafter referred to as junctions) into a single model, allowing for the detection of complex genomic rearrangements. We identified a median of 45 (range 0 - 212) simple structural variants and 8 (range 0 - 38) complex structural variants per sample (**Fig. 4a; Supplementary Tables S8 and S9**). 35/71 (49.2%) samples exhibited whole genome doubling (WGD), higher than the 20.4% reported in TCGA-UCEC and 37.8% in PCAWG^39^ respectively, but consistent with recent work showing an increased WGD rate in patients that self-report as B/AA.^40^

**Fig. 4:**
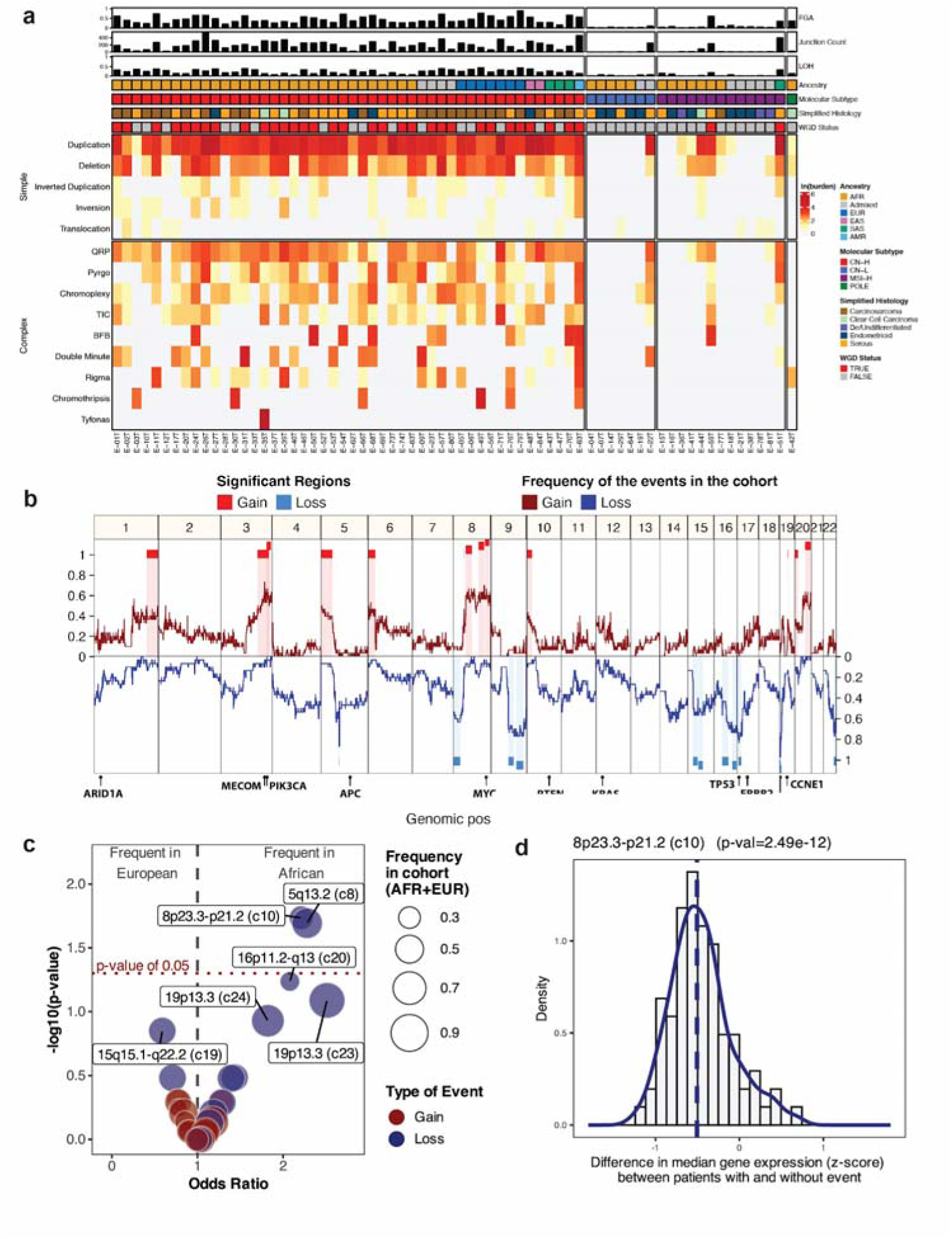
DNA related events and copy number alterations in the cohort with relevant ancestry associated events in the CN-H subtyping patients. **a,** Heatmap showing log-transformed junction burden of simple and complex structural variants (rows) in each tumor (columns). Samples are grouped by molecular subtype. Metadata annotations (from top to bottom) include fraction of genome altered (FGA), sample junction count, proportion of the genome with loss of heterozygosity (LOH), assigned ancestry, molecular subtype, simplified histology, and whole-genome doubling status. **b,** The frequency of the gain and loss events in the AFR patients assigned to the CN-H subtype annotated with the 30 recurrent large regions significantly detected. Genes of interest have been added to the lower section of the graph. **c,** The odds ratio for the frequency of the gain and loss events (30 regions) in the AFR and EUR patients assigned to the CN-H subtype in the cohort composed of TCGA-UCEC (WGS) and TCGA-UCS (WGS) and P-1000. The top regions are annotated. **d,** Distribution of the difference of the median gene expression (z-score variance stabilized transformed) for the genes present in the 8p23.3-p21.2 region identified inn **c**, between the patients with and without the detected event. All P-1000 patients assigned the CN-H subtype were retained for this analysis.

To ensure a comprehensive characterization of structural variation, we queried the cohort for the presence of extrachromosomal DNA (ecDNA). We identified 42 ecDNAs across 27 patients, 26 of which amplified known oncogenes (**Supplementary Table S10**). We observed recurrent ecDNA-mediated amplification of *FGFR3* and *NSD2* (n=2), and *MECOM* (n=2). *FGFR3* and *NSD2* reside approximately 60Kb from one another, and are co-amplified on the same ecDNA in both cases. 30/44 (68.2%) of ecDNA-amplified oncogenes were expressed in the top 5% of all samples (**Supplementary Fig. 11**).

CN-H tumors displayed a higher burden of structural variation as measured by fraction of genome altered (FGA), fraction of the genome with loss of heterozygosity (LOH), and total junction burden relative to CN-L tumors (p-value < 0.001, Wilcoxon rank sum test) (**Fig. 4a; Supplementary Fig. 12**), as well as a higher rate of whole genome doubling (WGD) (p-value<0.001, Fisher’s exact test). When limiting our analysis to the CN-H tumors, we observed a significantly higher fraction of the genome with LOH in carcinosarcomas when compared to serous tumors (p = 0.003, Wilcoxon rank sum test) and a depletion of templated insertion chains (TIC) in serous histology (p = 0.041, Fisher’s exact test). Within CN-H tumors, we did not detect any ancestry-specific associations of summary-level metrics for FGA, LOH, junction burden, and WGD, nor specific structural variant event classes.

We next sought to determine whether specific regions of the genome were differentially affected by large-scale and focal copy number alterations. We evaluated the presence of large regions with recurrent copy number events specific to our African patients (n=44). A total of 30 regions (13 recurring losses and 17 recurring gains) were significantly detected (**Fig. 4b; Supplementary Table S11**). All regions, except three, overlap at least one COSMIC annotated gene. Similar event frequencies were detected for African patients in both P-1000 (CN-H only, n=30) and TCGA-EC WGS data (TCGA-UCEC CN-H only and TCGA-UCS, n=25) (**Supplementary Fig. 13, Supplementary Table S12**).

The three most frequently deleted regions in the P-1000 CN-H African patients are in 19p13.3 (2 events identified as c23 and c24) and in 5q13.2 (identified as c8) with frequencies of 93.3%, 73.3%, and 86.6% compared to 88%, 80% and 56% in TCGA-EC. In the AFR CN-H patients, the 17p13.2-p11.2 region (identified as c22), which includes *TP53*, is recurrently lost (20/30 P-1000 and 11/25 TCGA-EC), as observed in many cancers^41^.

The 3q26.2 region (identified as c4) overlaps only four genes and is the most frequently observed gain event in the CN-H AFR patients (73.3% in P-1000 and 64% in TCGA-EC). This region includes the oncogene *MECOM* and is associated with poor outcomes in TCGA EC.^42^ This gain event is also detected at almost the same frequency (76%) in the TCGA-UCEC CN-H AFR patients. The 8q24.13-q24.3 gain region (identified as c14) was detected in 47% and 48% of the P-1000 and TCGA EC cohorts. This region overlaps the *MYC* oncogene. Frequent gain of the long arm of chromosome 8 (8q) has been observed in endometrioid endometrial carcinoma.^43^ Jiagge et al^44^ observed an enrichment of *MYC* gain in relation to African ancestry in non-small cell lung cancer, breast cancer, and prostate cancer. *MYC* gain was also associated with worse overall survival in those three cancers.

To augment our statistical power, we combined the P-1000 cohort with TCGA-UCEC and UCS WGS data to examine the differences in event frequency by ancestry (55 CN-H African vs. 119 CN-H European) across the 30 regions (**Fig. 4c; Supplementary Table S12**). Two regions were lost with significant ancestry-associated prevalence: 8p23.3-p21.2 and 5q13.2 (identified as c10 and c8) (p-value < 0.05, Fisher’s exact test). The genes in the 8p23.3-p21.2 region (c10) show decreased expression in tumors where the region is deleted (p<0.001, Wilcoxon rank test) (**Fig. 4d; Supplementary Table S13**).

### MECOM amplifications are enriched in patients of African ancestry

To identify regions with recurrent focal copy number alterations, we employed two complementary methods, one integrating junctions and adjusting for epigenomic covariates with a Gamma-Poisson model^45^ and the other using changes in copy number relative to ploidy.^46^ We noted coincident significant peaks in *MECOM*, *ESR1*, *MYC*, *SPOP*, and *IKZF3* (**Fig. 5a; Supplementary Tables S14** and **S15**). The prominent MECOM peak is contained within the c4 region previously identified in the analysis of large-scale copy number events. *MECOM* amplification was detected in 22/71 tumors (30.9%)(**Fig. 3a**), notably higher than TCGA-UCS (17.9%) and TCGA-UCEC (11.9%). These amplifications were especially frequent in patients of African ancestry (16/44 AFR, 2/4 SAS, 1/9 EUR, 1/1 AMR, 2/11 admixed, and 0/2 EAS patients).

**Fig. 5:**
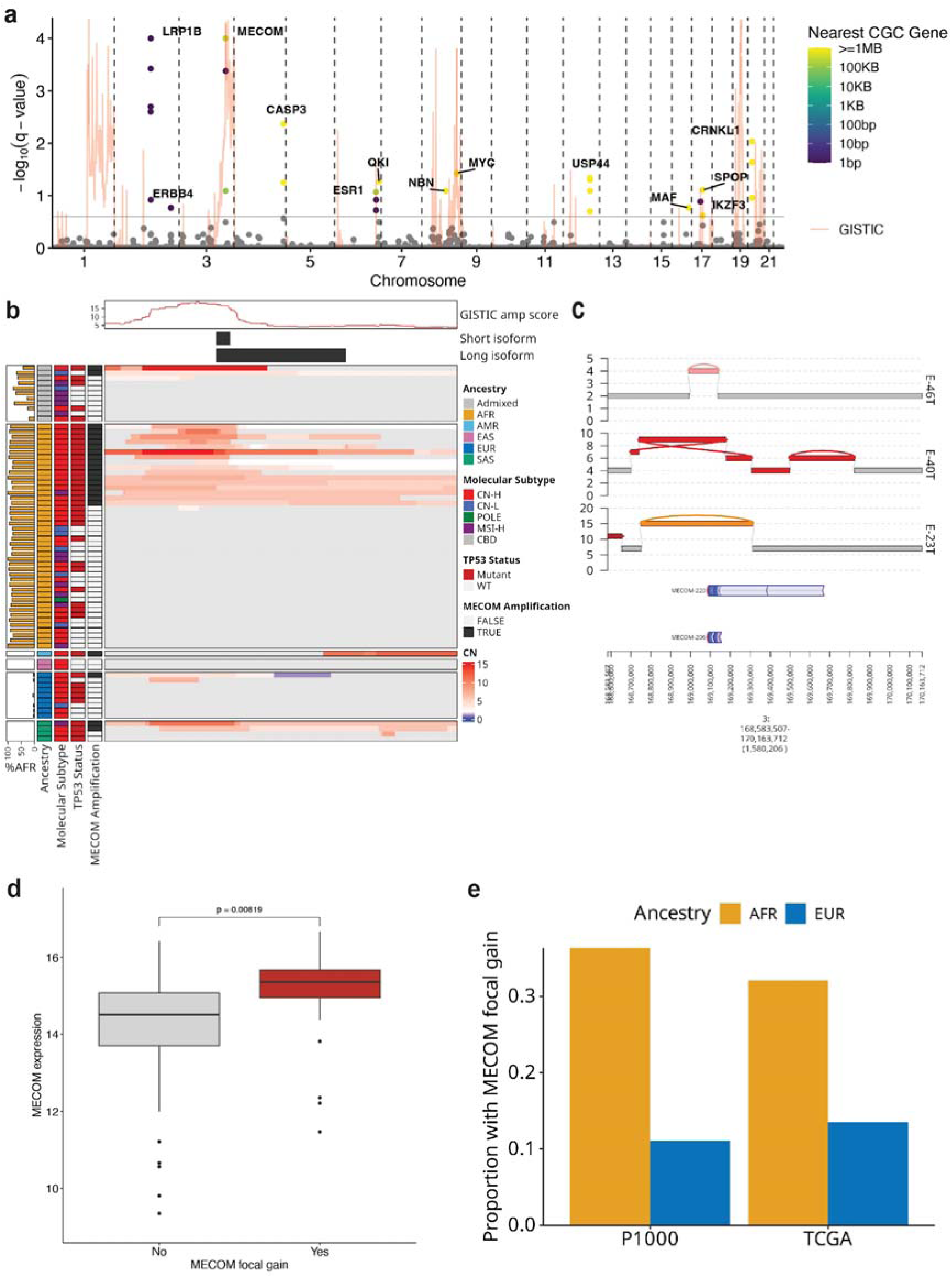
MECOM copy number gain is enriched in patients of African ancestry. **a,** Manhattan plot showing significance scores for the junction (FishHook) and copy-number (GISTIC) recurrence analyses. The red line indicates the amplification score as reported by the copy number analysis, and the points represent the junction recurrence scores of each 100KB window. The light gray horizontal line represents the FDR threshold used for both analyses. Points are labeled and colored by distance to the nearest Cancer Gene Census (CGC) gene, if the significance threshold is reached. **b,** Heatmap summarizing the landscape of focal copy changes at the MECOM locus. Segments longer than 3MB are masked in light gray. The GISTIC amplification score is reported in the top track, followed by a schematic representation of the canonical short (MECOM-206) and long (MECOM-220) isoforms. **c,** Example JaBbA plots showing different routes of amplification at the MECOM locus. The height of each bar indicates integer copy number, and the color of each bar represents membership in a structural variant (gray: none, pink: duplication, red: pyrgo, orange: double minute). The arcs connecting segments represent junctions and are colored in the same manner as segments. Thin gray arcs represent reference adjacencies, i.e. coordinates that are contiguous on the reference genome. **d,** Gene expression (batch-corrected, library-size-normalized, log-transformed), in samples with and without focal gain of MECOM. **e,** Barplot showing the proportion of samples with a MECOM focal gain stratified by ancestry in this cohort (P-1000), and in the whole-genome resequenced TCGA UCEC and UCS cohort.

*MECOM* (MDS1 And EVI1 COMplex Locus) is a master transcription factor^47,48^ previously implicated in leukemia,^49,50^ myelodysplastic syndrome,^51^ ovarian cancer,^52,53^ and more recently, EC.^42,54^ This minus-strand gene is under the control of two promoters separated by 531Kb and produces two transcriptional products, the 1042 residue protein EVI1, and the 1239 residue MDS1-EVI1 (alias PRDM3). Oncogenic activity has been ascribed to the short isoform EVI1, while expression of the long isoform MDS1-EVI1 is associated with a tumor suppressive effect.^55–58^ In this cohort, we noted that the short isoform was preferentially amplified (**Fig. 5b**), with a median copy number 1.39-fold greater than the long isoform-unique exons in MECOM-amplified cases (p < 0.001, Wilcoxon signed rank test). Surveying the landscape of complex structural variation around the locus, we found evidence of distinct amplification classes including tandem duplications, pyrgo, and ecDNA (**Fig. 5c**).

When comparing MECOM expression between samples with and without focal copy number gains, we observed a suggestive but not significant increase in expression (p = 0.0507, Wilcoxon rank sum test) (**Fig. 5d**). Expression was high in many copy-neutral cases, consistent with previous reports suggesting epigenetic mechanisms of upregulation.^54^ We did not identify any clear patterns distinguishing the short and long isoform of *MECOM* when looking at isoform-level estimations.

To confirm these findings, we reprocessed the recently released TCGA-UCEC and UCS WGS data through the same copy number pipeline. We noted a similar ancestry-specific effect, and when pooling the data and controlling for molecular subtype, we observed a significant enrichment of MECOM focal gains in AFR patients (33/97 vs 43/276 in EUR patients, p = 0.01, Mantel-Haenszel chi-squared test) (**Fig. 5e**). Furthermore, we confirmed a preferential amplification of the short isoform in the reprocessed TCGA data (**Supplementary Fig. 14**).

Prompted by a recent study reporting worse overall survival (OS) and progression-free survival in TCGA-UCEC CN-H patients with MECOM amplification,^42^ we tested the hypothesis that the focal amplification of *MECOM* confers a worse prognosis. Both large-scale CNV and focal amplification of *MECOM* were associated with worse overall survival (p<0.00045 and p<0.000037 respectively, Log-Rank test) and worse progression-free survival (p<0.00000009 and p<0.00000034 respectively, Log-Rank test) when compared with patients with no amplification of MECOM in TCGA. However, the two types of alterations are not statistically different from one another (p<0.477 for OS and p<0.702 for PFS) **(Supplementary Fig. 15, Supplementary Table S16)**.

Together, these data suggest that the focal amplification of the oncogenic isoform of MECOM is associated with poor outcome. This event is found to be more prevalent in patients of AFR ancestry, which could be related with the overall disparity we aimed to study. It also demonstrates the power of our whole-genome profiling approach to uncover a unique pattern of structural variation.

### Gene expression associations with African ancestry

Next, we leveraged bulk RNA-seq data from the same samples in order to identify associations between ancestry and gene expression. We focused our analysis on the largest subtype group in our cohort (CN-H, n=47). Three genes, *PLXNA3*, *ANKRD29* and *ADMTSL3,* showed significant associations with AFR ancestry proportion (Benjamini-Hochberg adjusted p-value < 0.10) (**Fig. 6a; Supplementary Table S17**). *PLXNA*3, a member of the plexin gene family, is a transmembrane semaphorin receptor involved in cytoskeleton remodeling and signal transduction and shows positive correlation with increasing AFR ancestry (**Fig. 6b**). *ANKRD29* and *ADAMTSL3* were negatively correlated with %AFR ancestry (**Fig. 6b**). Interestingly, *PLXNA3* and *ANKRD29* were identified in a pan-cancer analysis of ancestry-associated expression differences of the TCGA cohort, although a specific association in EC was not found.^28^

**Fig 6:**
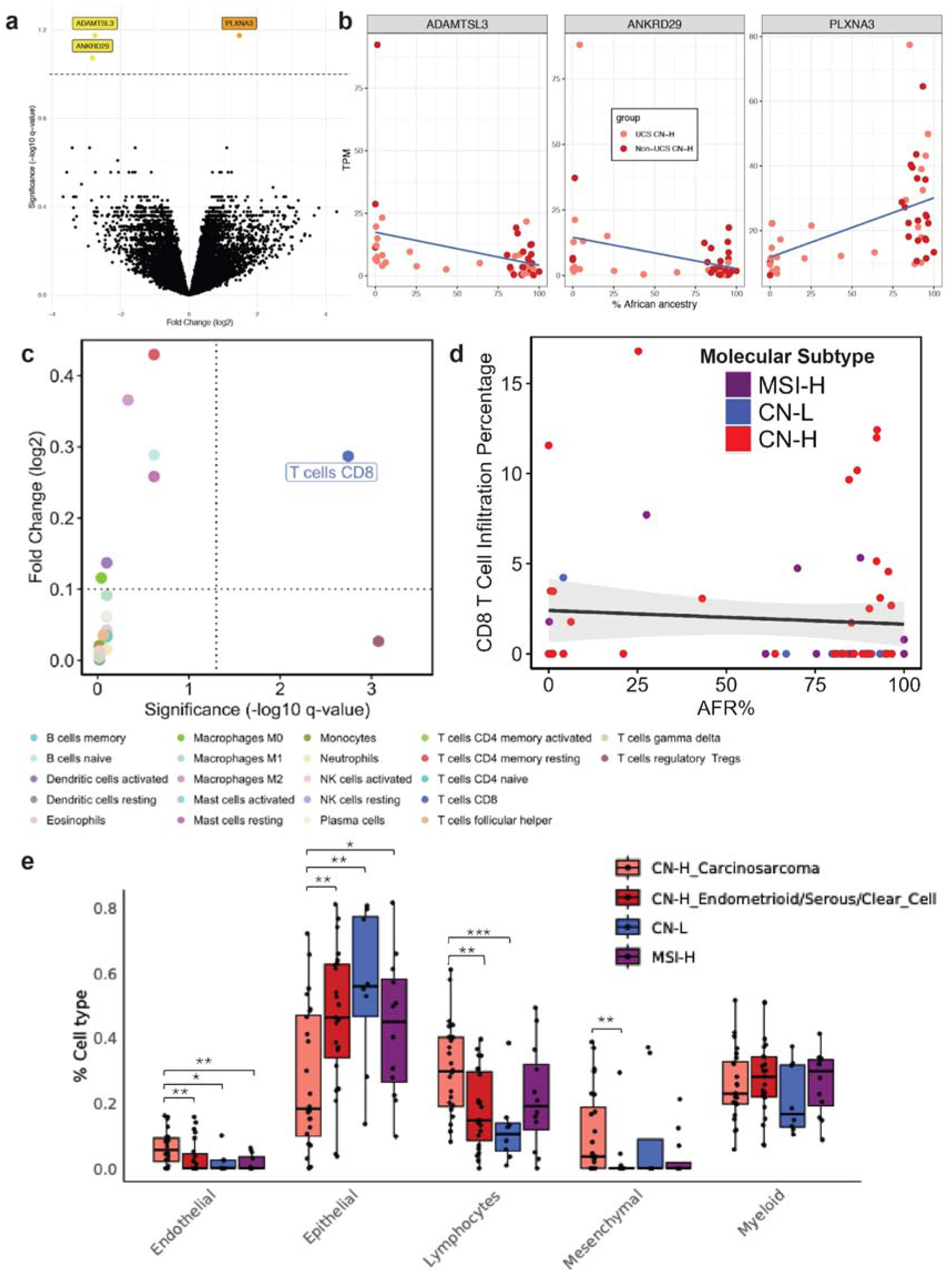
Ancestry-associated transcriptomic and immune landscape of high-grade endometrial cancer. **a,**Volcano plot visualizing the effect size (log2 Fold Change) and significance (-log10 Benjamini adjusted p-value) of the genes tested for ancestry-associated differential expression. Significant genes are highlighted in orange (positively correlated with African ancestry) and yellow (negatively correlated) **b,** Scatterplots depicting sample level expression levels by AFR ancestry ADMIXTURE coefficient for *ADAMTSL3*, *ANKRD29* and *PLXNA3*. The colors correspond to histological classification for carcinosarcoma (UCS) and non-carcinoma (non-UCS) CN-H samples. **c** Scatterplot visualizing the absolute effect size (absolute value of log2 Fold Change) versus significance (Benjamini-Hochberg adjusted p-value) for every infiltrating immune cell type for the comparison of interest (AFR% vs (EUR% + EAS% + SAS%), with dashed line indicating the significance thresholds (Benjamini-Hochberg adjusted p-value < 0.05 & absolute effect size ≥ 0.1). **d**, Estimated CD8 T Cell infiltration levels across samples with different African admixture levels. Samples are colored based on molecular subtype. **e**, Boxplot showing estimated lineage proportions across molecular subtypes with CN-H further split into carcinosarcoma and endometrioid/serous sub-groups. Statistical significance between comparisons are computed using the Wilcoxon rank sum test and only comparisons with significant differences are shown.

Gene set enrichment analysis (GSEA) showed that “MYC targets” and “DNA repair” hallmark genesets were enriched in genes positively correlated with African ancestry in the CN-H cohort (**Supplementary Fig. 16a; Supplementary Table S18**). Hallmark genesets enriched in genes negatively correlated with African ancestry include “androgen response”, “fatty acid metabolism”, and “adipocyte development”. Pathway activation analysis identified significant positive associations between African ancestry proportion and activation of the hypoxia and NF-κB pathways **(Supplementary Fig. 16b)**. In contrast, the TGF-β, TRAIL (TNF-related apoptosis-inducing ligand), estrogen, androgen, p53, and JAK–STAT pathways exhibited negative activation scores with increasing African ancestry. These pathway enrichment in genes associated with African ancestry replicated broadly in the TCGA cohort (**Supplementary Fig. 17; Supplementary Table S19**) and were corroborated by GSEA results on Gene Ontology Biological Process (GO BP) (**Supplementary Fig. 18; Supplementary Tables S20 and S21**).

#### Ancestry correlations with tumor infiltrating immune cells

To investigate how immune infiltration varies with ancestry, we estimated the relative proportion of 22 immune cell types using bulk gene expression data **(Supplementary Fig. 19, Supplementary Table S22)**. While both CD8 T cells and T regulatory cells showed significant differences in infiltration by ancestry, only CD8 T cells demonstrated a biologically meaningful effect size (≥ 0.1) **(Fig. 6c)**. Specifically, the analysis revealed decreased CD8 T cell infiltration with an increasing proportion of African admixture **(Fig. 6d)**. This finding is particularly compelling given the essential role of CD8 T cells in anticancer immune responses and the improved prognosis associated with higher infiltration of active CD8 T cells in solid tumors.^59^

#### Cellular lineage deconvolution via a diverse single-cell reference

To investigate the cellular lineage composition of tumors in this cohort, we used endometrial single-cell transcriptomic reference atlas data generated from a diverse cohort of non-disease donors (manuscript in preparation) to deconvolute bulk tumor expression profiles.

When considered independently, molecular subtypes showed minimal significant difference in estimated lineage proportions (**Supplementary Fig. 20a; Supplementary Table S23**). In contrast, jointly stratifying by molecular subtype and tumor histology revealed clear distinctions, most notably between CN-H carcinosarcoma and all other subtype-histology groups. CN-H carcinosarcoma samples had higher endothelial, lymphocytes and mesenchymal and lower epithelial lineage proportions relative to other molecular groups (**Fig. 6e**). Within the CN-H subtype, histological variability was also evident as carcinosarcoma samples displayed higher mesenchymal and endothelial lineage proportions relative to CN-H endometrioid/serous tumors (**Fig. 6e**), consistent with the biphasic nature of carcinosarcoma.^60^ This pattern was reinforced when estimating epithelial-to-mesenchymal (E/M) ratios (**Supplementary Fig. 20b**) and hallmark epithelial mesenchymal EMT scores **(Supplementary Fig. 20c)**. The same result was found when using the published Human Endometrial Cell Atlas as reference^61^ **(Supplementary Fig. 21).** We found no significant associations between the estimated cell lineage proportions and proportion of AFR ancestry (all p > 0.05; **Supplementary Fig. 22**) in models that adjusted for tumor purity and molecular subtype, revealing that cellular lineage composition was shaped primarily by histological and molecular characteristics of the tumors.

## Discussion

Endometrial cancer is characterized by profound heterogeneity across its histological subtypes and molecular landscapes. This intrinsic biological complexity is further compounded by persistent racial and ethnic disparities in EC incidence, tumor aggressiveness, and patient outcomes, particularly impacting women of African ancestry. Despite significant advances in genomic characterization, the biological contributions of genetic ancestry to these complex molecular profiles, hypothesized to underpin a portion of these disparities, have remained largely underexplored. This study aimed to elucidate the impact of genetic ancestry on the genetic and molecular drivers of EC heterogeneity, thereby offering crucial insights into the biological determinants of observed racial and ethnic disparities in EC. This comprehensive WGS approach allowed for in-depth characterization of multiple mutation types, including SNVs, indels, copy-number variants and structural variants, across both coding and non-coding regions, providing a new understanding of the disease’s genomic architecture.

Unlike many prior EC studies, our recruitment strategy specifically targeted a diverse patient population, with a particular focus on more aggressive, high-grade tumors that have the worst prognosis. This deliberate enrichment of individuals of AFR ancestry proved instrumental, providing a unique lens through which to interrogate the understudied role of genetic ancestry in EC heterogeneity. The somatic mutational profiles largely recapitulated genes previously implicated in EC carcinogenesis, affirming the fundamental genomic alterations common across the disease. However, striking findings emerged from ancestry-stratified analyses. We identified several putative cancer driver genes such as *ZBTB7B* and *ARHGAP35* exclusively mutated in patients of AFR ancestry. While the relatively low number of observations for these specific mutations in our cohort necessitates further validation, these genes represent intriguing candidates for ancestry-specific tumorigenesis and warrant deeper investigation into their functional roles and potential therapeutic vulnerabilities in AFR-ancestry EC.

Beyond SNVs, the evaluation of large-scale CNVs highlighted two regions (8p23.3-p21.2 and 5q13.2,) with recurrent losses, more prevalent in the AFR-ancestry patients. Somatic recurrent deletion of 8p has already been described in EC^62^ and more specifically in endometrial serous carcinoma.^63^ However, no previous ancestry-associated prevalence has been described for those two regions in EC.

Our analysis also demonstrated the critical importance of comprehensively evaluating both large-scale and focal copy-number alterations using WGS. Although the amplification of *MECOM* was recently associated with an aggressive form of EC^42^, our study further demonstrated that the focal amplification of the oncogenic isoform of *MECOM* was enriched in patients of African ancestry and was likely a stronger contributor to poor prognosis than the large-scale amplification of the gene. This finding suggests that distinct molecular pathways may contribute to EC pathogenesis in different ancestral populations. Further insights into ancestry-associated molecular differences include the observation of CN-H tumors predominantly in patients of AFR ancestry, notably independent of TP53 mutation status. This suggests alternative mechanisms driving genomic instability in this subgroup. Concurrently, we found an association between African ancestry and decreased CD8+ T-cell infiltration, hinting at potential ancestry-specific immune microenvironment characteristics that could influence disease progression and expand therapeutic options.

Despite being the largest WGS study of EC to date and offering unprecedented ancestral diversity, the principal limitation of our study remains its overall cohort size. This limited our statistical power to robustly identify and validate novel ancestry-specific driver genes, particularly given the significant histological and molecular heterogeneity intrinsic to EC. This variability also precluded crucial subgroup analyses that would have enabled more granular characterization of ancestry-specific effects and their effects on clinical outcomes. While we compared our findings with publicly available cohorts, some novel observations lack independent validation, in part due to inherent biological and methodological differences across disparate study cohorts. Therefore, larger, well-characterized studies with broader ancestral representation are needed to confirm these preliminary findings.

In summary, this study reveals the value of ancestrally diverse cohorts for understanding the interplay between genetic ancestry and the molecular drivers of cancer heterogeneity. Ancestry emerges, not simply as a demographic variable but as a biological determinant that shapes the genomic landscape of EC, affecting driver genes, copy-number alterations, and the tumor microenvironment. These findings highlight the necessity for inclusive research designs for equitable and effective precision oncology.

## Methods

### Patient Recruitment/IRB Study Approval

The study was IRB approved (#18-0897) at Northwell Health. Patients were screened based on preoperative diagnosis of endometrial cancer - low grade and high grade. Patients undergoing hysterectomy for endometrial cancer were recruited at time of surgery at Northwell Health. Eligible patients included women >18 years old diagnosed with endometrial adenocarcinoma, uterine serous carcinoma, carcinosarcoma, dedifferentiated carcinoma, undifferentiated carcinoma, or clear cell carcinoma. Women who had received prior treatment, either chemotherapy or radiation therapy were excluded. Patients signed study specific informed consent prior to surgery.

### Sample/Clinical Data Collection

Endometrial cancer tissue, benign endometrium, and blood were collected on all patients at time of hysterectomy. Tissue was taken to the pathology lab where specimens were collected. Each participant had an assigned study ID number and all specimens were transferred fresh to the laboratory. The lab staff were blinded to subject and treatment status. Once delivered to the laboratory, the samples were assigned laboratory IDs and stored at -20 degrees C prior to processing. For all recruited patients, demographic and clinical data was collected including: race, ethnicity, tumor histology, recurrence and survival data, pathology reports and molecular subtype testing.

#### Snap Freezing

Fresh benign or malignant tissue samples were washed with cold Ca²L/Mg²L-free 1X PBS (homemade), minced into 3–4 cubic millimeter sections, and snap frozen in liquid nitrogen. The samples were then transferred to a –80°C freezer for long-term storage.

#### PBMC Isolation

PBMCs were isolated from patient whole blood using Lymphoprep (Stemcell Technologies, 18060) and SepMate-50 tubes (Stemcell Technologies, 85450) following the manufacturer’s instructions. The isolated PBMCs were cryopreserved following resuspension in Recovery Cell Culture Freezing Medium (ThermoFisher, 12648010).

#### DNA/RNA Extraction

DNA and RNA extraction was primarily conducted at NYGC using the following protocol: PBMC normal samples were extracted with the QIAamp DNA Blood Mini Kit (Qiagen, 51106). Frozen tissue samples were extracted with Qiagen’s AllPrep kit (80204) if both DNA and RNA were needed, the QIAamp DNA Mini Kit (Qiagen, 51306) if only DNA was needed, or the RNeasy Mini kit (Qiagen, 74106) if only RNA was needed. For all kits the manufacturer’s instructions for input and protocol were followed. For a subset of samples, extractions were performed at CSHL using the following protocol: DNA was extracted from PBMCs isolated from patient blood or snap frozen tissue using the Zymo Quick-DNA Miniprep kit (Zymo, D3024) following the manufacturer’s instructions. DNA quality and concentration were measured using a Nanodrop ND-1000 Spectrophotometer. Total RNA was extracted from snap frozen tumor tissue using the Zymo Quick-RNA Miniprep kit (Zymo, R1054) with DNase treatment steps, following the manufacturer’s instructions. RNA quality and concentration were measured using a Nanodrop ND-1000 Spectrophotometer. There was no difference in DNA yield and quality based on the extraction site (**Supplementary Table S1**).

### Whole Genome Sequencing

#### Library preparation

Whole genome sequencing (WGS) libraries were prepared using the Truseq DNA PCR-free Library Preparation Kit (Illumina) in accordance with the manufacturer’s instructions. Briefly, 1 ug of DNA was sheared using a Covaris LE220 sonicator (adaptive focused acoustics). DNA fragments underwent bead-based size selection and were subsequently end-repaired, adenylated, and ligated to Illumina sequencing adapters. Final libraries were quantified using the Qubit Fluorometer (Life Technologies) or Spectromax M2 (Molecular Devices) and Fragment Analyzer (Advanced Analytical) or Agilent 2100 BioAnalyzer. Libraries were sequenced on an Illumina Novaseq6000 sequencer using 2x150bp cycles.

#### WGS Data Pre-processing

The New York Genome Center somatic pipeline (v6)^64^ was used to process and align the WGS data and call variants. Sequencing reads for the tumor and normal samples were aligned to reference genome GRCh38 using BWA-MEM (v0.7.15)^65^. Short reads were marked as unaligned and removed with NYGC’s ShortAlignmentMarking tool (v2.1)^66^. GATK (v4.1.0)^67^ FixMateInformation was run to verify and fix mate-pair information, followed by Novosort (v1.03.01), markDuplicates to merge individual lane BAM files into a single BAM file per sample, coordinate sort and marked duplicated. GATK’s base quality score recalibration (BQSR) was performed.

#### Variant Calling and Filtering

MuTect2 (GATK v4.0.5.1),^68^ Strelka2 (v2.9.3),^68^ and Lancet (v1.0.7)^69^ for calling SNVs and indels, Svaba (v.0.2.1)^70^ for calling indels and SVs, and Manta^71^ (v1.4.0) and Lumpy (v0.2.13)^72^ for calling SVs.^72^ The candidate set of indels output from Manta was used as input to Strelka2 as per the developer’s recommendation. Germline SNPs and indels were called on the matched normal samples with GATK HaplotypeCaller (v3.5) and filtered with GATK VQSR at tranche 99.6%. The positions of heterozygous germline variants were used to compute B-allele frequencies (BAF) in the tumor samples. Variant calls were merged by variant type (SNVs, indels, and SVs) and annotated using the following databases: Ensembl^73^ (v93)^73^, COSMIC (v86),^36^ 1000Genomes (Phase 3),^74^ ClinVar (201706),^20^ PolyPhen (v2.2.2), ^75^SIFT (v5.2.2),^76^ FATHMM (v2.1),^77^ gnomAD (r2.0.1),^78^ and dbSNP (v150)^79^ using Variant Effect Predictor (v93.2)^80^. Actionable mutations were annotated with OncoKB MafAnnotator (v3.4)^81^, with the tumor type set to UCEC and considered with regards to Therapeutic Levels of Evidence V2. Somatic variants that occurred in two or more individuals in an in-house panel of normals, SNV/indels that had minor allele frequency >= 1% in 1000Genomes or gnomAD, and SVs overlapping with DGV (2020-02-25 release),^82^ 1000Genomes or gnomAD-SV (v2.0.1)^83^ were removed. SNV/indels with tumor VAF < 0.0001, normal VAF > 0.2, or depth < 2 in either the tumor or normal sample, or normal VAF greater than tumor VAF were filtered from the final callset. SNV/indels in the final callset were marked as high confidence if there was support from at least two callers. SVs with support from at least two callers or one caller with additional support from a nearby BIC-Seq2 (v0.2.6)^84^ CNV changepoint or split-read support from SplazerS^85^ were marked as high confidence.

#### Concordance and contamination assessment

We ran Conpair^86^ to confirm that paired tumor and normal samples were derived from the same patient, and to estimate any inter-individual contamination. DeTiN^87^ was used to estimate tumor-in-normal contamination using Mutect2 calls marked PASS or “normal_artifact” with NLOD greater than -25, read depth, and BAF as input.

#### Genetic Ancestry Inference

We inferred genetic ancestry proportions relative to reference populations from the 1000Genomes project using ADMIXTURE (v1.3.0).^88^ The software uses a maximum likelihood-based method to estimate the proportion of reference population ancestries in a given sample. We genotyped the reference markers generated from 1,964 unrelated 1000Genomes project^89^ samples directly on the study samples using GATK pileup. Individuals from the MXL (Mexican Ancestry from Los Angeles USA), ACB (African Caribbean in Barbados), and ASW (African Ancestry in Southwest US) populations were excluded from the reference set due to known high admixture. The reference was further filtered by using only SNP markers with a minimum minor allele frequency (MAF) of 0.01 overall and 0.05 in at least one 1000Genomes superpopulation. Variants are additionally linkage disequilibrium (LD) pruned using PLINK v1.9 with a window size of 500kb, a step size of 250kb and r2 threshold of 0.2. The analysis results in a proportional breakdown of each sample into 5 continental populations (AFR, AMR, EAS, EUR, SAS) and 23 subcontinental populations.

The continental and sub-continental ancestral admixtures of The Cancer Genome Atlas (TCGA) patients (RRID:SCR_003193) with multiple types of cancers were inferred using an in-house version of RAIDS^90^ software (RRID:SCR_027265) invoking ADMIXTURE software (v1.3.0)^88^ in supervised mode (RRID:SCR_001263) as a component. The final results include the proportional breakdown into 5 continental and 18 subcontinental populations for each TCGA patient.

The ancestry distribution plot was generated using the Bioconductor ComplexHeatmap package (v2.22.0).^91^ The ancestry to race graph was generated using the CRAN networkD3 package (v0.4)^92^. The distribution of the African related subcontinental ancestry in the P-1000 and TCGA cohort was generated with CRAN ggplot2 package (v4.0.0)^93^(RRID:SCR_014601).

#### Germline Pathogenic Variant Prioritization

Germline variants for each individual were annotated with snpEff (v4.3, GRCh38.82)^94^ and AnnoVar (Jun 2020, hg38)^95^ using the following databases; refgene, COSMIC (v98), dbNSFP47a, GnomAD4 genomes and exomes, and Revel. Further, an internal modified version of ClinVar (sept, 2024) was used to annotate the variants. ClinVar modification was done by calling conflicting variants based on the following criteria: 1) based on the majority of calls, 2) in-case no majority was identified, based on the most recent call.

Variants of interest were identified by an internally developed scoring scheme based on ACMG guidelines including PVS1, PS1, PM2, PP2, PP3 and PP5 for pathogenic criteria and BA1, BP1, BP4 and BP7 for benign criteria.^96^ Each criterion was assessed as either having very strong, strong, moderate, or supportive evidence with a corresponding 8, 4, 2 or 1 points for those that support pathogenicity and -8, -4, -2 or -1 point for criteria that support a benign outcome. Variants of unknown significance were given a score of 0. The final scores > 8 points are classified as likely pathogenic and > 11 points as pathogenic (P/LP). Germline P/LP variants in a curated cancer predisposition gene set (CPG) or within DNA repair genes (**Supplementary Table S4**) were prioritized.

#### Somatic variant prioritization and cohort mutational frequency comparison

Variant calls from all samples were annotated with ANNOVAR ^95^ (RRID:SCR_012821) using a broad range of variant assessment tools including prediction of deleteriousness (dbNSFP v41a (RRID:SCR_005178), SIFT^97^ (RRID:SCR_012813), Polyphen^75,98^ (RRID:SCR_013189), MutationTaster^99^ (RRID:SCR_010777), etc) and conservation scores (CADD ^100^ (RRID:SCR_018393), GERP (RRID:SCR_000563), DANN^100,101^, etc). We selected rare loss of function variants (nonsense, frameshift, splice site) with frequency less than 1% in the gnomAD v4^102^ (RRID:SCR_014964), ExAc^103^ (RRID:SCR_004068), and 1000 Genomes (RRID:SCR_006828) databases. Missense and in-frame indel variants were selected if they were noted as pathogenic by ClinVar 20250721^104^ (RRID:SCR_006169), or if they are are both rare and annotated as pathogenic by COSMIC v96^36,103^ (RRID:SCR_002260), or if they are both rare and found to be present in the TCGA^105^ (RRID:SCR_003193) or ICGC^106^ (RRID:SCR_021722) cohorts.

TCGA data were downloaded from TCGA GDC^107^ and MSKCC data were downloaded from cBioPortal^108^ Further evaluation of these candidate variants was performed using Maftools v2.14 ^109^ (RRID:SCR_024519). Variant frequency per gene across samples was assessed and variant summaries and oncoplots were generated. AFR vs EUR cohorts were compared using the Maftools function mafCompare, performing a Fisher’s exact test to assess differentially mutated genes in each sample set. Variants had to present in a minimum of 3 samples per cohort to avoid bias of mutations in a single sample. Additional analysis was performed with maftools `oncopathways`.

### Detection of genes under positive selection

Detection of cancer driver genes was done with the dNdScv R package (v0.1.0)^30^ using default parameters and the recommended resource files for GRCh38. cBioPortal MutationMapper^110^ was used to visualize and further analyze the mutations associated with driver gene FBXW7.

#### Mutational signatures

Mutational signature fitting to COSMIC (v3.4)^36^ was performed using the python package MuSiCal (v1.0.0) on the SNV/indel high confidence callset with the refit function with method set to “likelihood_bidirectional” and thresh of 0.001.

#### Purity and ploidy estimation

Purity and ploidy were estimated for each tumor-normal pair using AscatNGS (v4.2.1)^111^ and Sequenza (v3.0.0).^112^ Estimates were manually reviewed and chosen based on fit to observed VAF, BAF and read depth. All estimates were subjected to automated QC with CNAqc (v1.1.0)^113^ to confirm validity.

#### Complex structural variation and copy number calling

Tumor read depth was collected in 1KB bins and corrected for genomic GC content and mappability using fragCounter^114^. Corrected tumor coverage profiles, BAF, purity/ploidy estimates, and high confidence SVs were used as input to JaBbA (v1.1);^38^ default parameters were used, with the exception of `rescue.all` set to false, `maxna` set to 0.8, `slack` of 1000 and `ism` set to true. The JaBbA companion R package gGnome (commit c390d80)^38^ was used to call simple inversions, translocations, duplications and deletions, inverted duplications, chromoplexy, chromothripsis TICs, quasi-reciprocal pairs, rigma, pyrgo, tyfonas, breakage fusion bridge cycles and double minutes on the junction balanced genome graph. Using the JaBbA output integer copy number, the fraction of genome altered (FGA) was computed as the proportion of autosomes not in a neutral copy state (defined by sample ploidy). Samples with an intermediate average ploidy (fractional value of 0.4-0.6, for example, 3.5), the copy-neutral state was set as the closest two integer values, otherwise the copy-neutral state was set as the rounded ploidy.

The TCGA UCEC and UCS data were processed in a similar manner, with the exceptions that the purity and ploidy values were obtained from the PanCanAtlas publication page^115^, the breakpoint calls were obtained from the GDC^107^, and the slack penalty was set to 50, as it was noted that many true copy changes were missing accompanying breakpoints. Tumors without complete purity, ploidy, and breakpoint calls were excluded, as were those without molecular subtyping.

AmpliconSuite-pipeline (v1.15.2)^116^ was run with default parameters on tumor BAMs using JaBbA-inferred total copy number to derive seed intervals. Only intervals with total copy number greater than 4 and longer than 10KB were considered.

Recurrent focal amplifications and deletions were identified with GISTIC2 (2.0.23),^46^ using default parameters. The log_2_ of total copy number normalized by sample ploidy was used as input, with a cutoff of 0.58 for amplifications and -1, equivalent to a single copy change in either direction for a diploid sample.

Recurrent regions of structural variation were detected with FishHook^45^. 100kb non-overlapping windows were generated across the genome, with regions of poor mappability excluded using a coverage mask described in ref ^117^. Junctions incorporated into the JaBbA junction-balanced genome graphs were used as input. To avoid bias towards complex events that are present in few samples, a given sample was allowed to contribute at most one junction in a given bin. To adjust for the background mutation rate, replication timing^118^, fragile sites^119^, di- and trinucleotide frequency, RepeatMasker LINE, SINE, simple repeat, and transposon elements^120,121^ were included as covariates in the model. An FDR-adjusted p-value of 0.25 was used as a significance threshold.

#### Large regions with recurring gain and loss events in the African assigned patients

Recurrent gain and loss events were identified, using the P-1000 African patients (n=44), with CRAN CORE package v3.2^122^ (RRID:SCR_027419) using the Core I setting for larger event detection with R software v4.4.0 (RRID:SCR_001905). The copy numbers obtained from JaBbA were used as input, excluding the HLA region (chr6:28600000-33200000) and the blacklisted regions (**Supplementary Table S15**). A total of 30 regions were significantly detected (**Supplementary Table S11**). The frequency of the events in the P-1000 patients assigned with CN-H tumors was calculated and the associated frequency plot was generated with Bioconductor gtrellis package v1.38.0^123^ (RRID:SCR_027267).

Genes overlapping those regions were identified using the Bioconductor TxDb.Hsapiens.UCSC.hg38.knownGene v3.20.0 and GenomicRanges^124^ (RRID:SCR_000025) v1.58.0 packages. Among those genes, the one associated with cancer were identified using the COSMIC Cancer Gene Census database v102^125^ (RRID:SCR_002260).

The gain and loss copy-number values for the 30 recurrent regions were extracted from TCGA-EC WGS patients^9,126^. Patients with African admixture, as defined in ^28^ with an admixture level of 80% or higher were assigned to the African cohort. The event frequency for each region was calculated for the TCGA-EC cohort composed of the TCGA-UCEC African patients assigned to the CN-H subtype and all the TCGA-UCS AFR patients (**Supplementary Table S11**).

Using the African and European patients assigned to the CN-H subtype for the combined P-1000 and TCGA-EC cohorts (all TCGA-UCS African patients included), the difference in event frequencies for each core was tested using a Fisher’s Exact test (**Supplementary Table S12**). The associated graph was generated with the CRAN ggplot2 v4.0.0^93^ (RRID:SCR_014601).

For each region, the difference in the gene expression between the P-1000 patients assigned to the CN-H subtype (n=47) with and without the event has been tested. A Wilcoxon rank test was performed on the difference in median expression (z-score) using each gene present in the region (**Supplementary Table S13**). The associated graph was generated with the CRAN ggplot2 v4.0.0^93^ (RRID:SCR_014601).

#### Survival analysis

Overall survival, progression-free survival, and disease-free survival of TCGA-UCEC and UCS cohorts was performed using the R survminer package ^127^. Pairwise comparisons were performed with the Log-Rank test using the pairwise_survdiff function with default parameters.

#### MSI Detection

MANTIS^128^ (v1.0.4) was run for Microsatellite Instability (MSI) detection in microsatellite loci (found using RepeatFinder, a tool included with MANTIS). A sample is considered to be 6 microsatellite unstable if its Step-Wise Difference score reported by MANTIS is greater than 0.4 (or 0.62 in absence of a matched-normal). Otherwise it is considered to be microsatellite stable3 (MSS).

#### Molecular Subtyping of Cohort

We performed molecular subtyping according to the four subtype scheme identified by the Cancer Genome Atlas (TCGA)^9^: POLE mutants (POLE), Microsatellite Instability High (MSI-H), Copy Number Low (CN-L), and Copy Number High (CN-H). Following the methodology described in^9^, we classified POLE mutant samples based on somatic hotspot mutations in the POLE exonuclease domain. MSI-H cases were determined using mismatch repair (MMR) pathway deficiency immunohistochemistry (IHC) staining as a part of routine clinical care. For patients without MMR IHC data, MSIsensor^129^ and MANTIS^128^ were used to infer MSI status from paired tumor normal whole genome sequencing. Patients that were MMR deficient and/or who had above the 10 and 0.4 threshold for MSIsensor and MANTIS respectively were classified as MSI-H. CN-L and CN-H were determined using the method previously described by the Clinical Proteomic Tumor Analysis Consortium (CPTAC).^16^ Tumors who had more than 10% of their genome deleted were classified as CN-H and those with less than 10% of their genome deleted were classified as CN-L. Samples with the histology “Other” were excluded from molecular subtyping as there is limited data on the prognostic value of the subtypes within these diseases.

### RNA Sequencing

#### RNA Library Preparation

RNA libraries were prepared using the KAPA Stranded RNA-seq with RiboErase (H/M/R) library preparation kit (Roche 07962304001) in accordance with the manufacturer’s instructions. 500ng of total RNA samples were ribodepleted using oligonucleotide hybridization and RNase H treatment followed by DNase treatment. The RNA was fragmented using divalent cations under elevated temperature. The cleaved RNA fragments were copied into cDNA complementary molecules, adenylated, ligated to Illumina sequencing adapters, purified and enriched with PCR to create the final cDNA library. Final libraries were quantified using the Qubit Fluorometer (Life Technologies) or Spectramax M2 (Molecular Devices) and evaluated for size distribution on the Fragment Analyzer (Agilent). Libraries were sequenced on an Illumina Novaseq6000 sequencer using 2x100bp cycles.

#### RNA-Seq Data Processing and Analysis

Reads were aligned using the STAR aligner (v2.5.2a),^130^ versus the GRCh38 genome subsetted to only canonical chromosomes (chr1-22, X, Y, and M), with Gencode v25 as the annotation. Genes were quantified using featureCounts from the subread package (v1.4.3-p1).^131^ Quality was evaluated using Picard CollectRnaSeqMetrics.^132^ DESeq2^133^ was used to normalize gene-level counts across samples. FusionCatcher^134^ was used for fusion detection. Conpair^86^ was used to confirm sample identity versus matched DNA. One sample (E-76T) was found to be discordant with DNA, and so was removed.

For the initial RNA analysis, all 82 samples passing RNA QC and concordant with tumor DNA were included (including those that failed DNA QC or with high tumor in normal contamination, metastases, and "other" histology). E-46T RNA was dropped prior to sequencing, due to poor quality.

PCA analysis was performed on log-transformed, library-size-normalized counts using the "pca" module from the PCAtools package^135^ with removeVar = 0.1. An eigencor plot revealed that PCs 2 and 3 were strongly associated with the sequencing batch, with the first batch (B01) strongly separating from the subsequent two (B02 and B03) (**Supplementary Fig. 23**).

To create a corrected count matrix, batch correction was performed on the counts data using the ComBat-Seq method^136^ in R (version 4.4.1) via the sva package (version 3.54.0). B02 and B03 were combined as one batch. Genes with zero counts in more than 20% of samples were excluded. The argument covar_mod was used to retain the effect of continental genetic ancestry (design ∼AFR+SAS+EAS+AMR).

For downstream analysis, these batch-corrected counts were then subsetted to remove any samples that were excluded from the DNA analysis, leading to n=69 remaining RNA samples. PCA analysis was then performed again on the corrected matrix, using the same methods as before correction.

For the subsequent eigencorplot, correlations were calculated separately for each molecular subtype (CN-H, CN-L, and MSI-H) versus the other two, and same for histology (carcinosarcoma, serous, and endometrioid) (**Supplementary Fig. 24**).

#### Gene Expression Associations with Ancestry

To identify gene expression patterns associated with AFR ancestry across the CN-H samples, we created a multivariate linear regression model (*limma* R package version 3.62.2^137^), using tumor purity and histology - carcinosarcoma (UCS) versus all others (non-UCS), as additional covariates. African ancestry was treated as a continuous variable. The batch-corrected counts from ComBat-Seq, subsetted to only protein coding genes, were used as input. Then, calcNormFactors was used for library size normalization. Then, the modules voom, lmFit, eBayes, and topTable were used to create a results table.

TPM (transcripts per million) was calculated using the union of all possible exons as the gene length. The set of genes used was the same as what was input into differential expression (coding genes only). For the TCGA analysis, STAR counts for the UCEC and UCS cohorts were downloaded from GDC. ComBat-seq was once again used for batch correction, with an expression filter of genes having no more than 20% zeroes. Metadata was obtained from Carrot-Zhang et al.^9^, as well as molecular subtype information for UCEC from Sanchez-Vega, Mina, and Armenia et al.^126^. Next, platform (IlluminaGA versus IlluminaHiSeq) was used as the batch variable, with cohort (UCEC versus UCS) as the “group” variable (for which we wanted to retain variation). After that, both cohorts were subsetted to samples with tumor purity and ancestry information available, and for the UCEC cohort, with molecular subtype information available. For differential expression, these batch-corrected counts, subsetted to protein coding genes, were then used as input into limma. The UCEC cohort was subsetted to CN-H only (resulting in n=153 UCEC samples), while all UCS samples were retained (since molecular subtypes were not available). The design was AFR + purity + histology (UCEC vs. UCS).

The “t” column (which is signed, and higher magnitude for more significant genes) from the limma output was used to rank genes for input into gene set enrichment analysis (GSEA). WebGestalt^138^ was used, with a cutoff of FDR 5% for calling gene sets as significant. For the GO BP gene sets, weighted set cover was used for redundancy reduction to select top categories for display, with “number of categories expected from set cover” set to 10.

#### Immune Deconvolution Analysis

To investigate infiltrating immune cell proportions across primary tumor samples we ran CIBERSORT utilizing the software’s LM22 immune cell gene expression signature matrix as reference matrix. Cell subset infiltration scores were obtained as fractions.

To further examine infiltration immune cell differences by ancestry we ran a multivariate linear regression model, while controlling for molecular subtype and this time comparing fractions of 22 immune cell type as the proportion of AFR ancestry (AFR%) compared to the proportions of EUR, EAS and SAS ancestries varied across the patients (AFR% vs (EUR% + EAS% + SAS%)). Proportion of AMR admixture (AMR%) was not included in the model since the AMR admixture levels were very low across all samples. After exclusion of a POLE-mutant and an MSI-H carcinosarcoma sample, the remaining (n=67) RNA-seq samples that passed QC were included in this analysis. Significantly differentially tumor infiltrating immune cell populations were identified at Benjamini-Hochberg^139^ adjusted p-value 0.05 and effect size ≥ 0.1.

#### Pathway Activation Analysis

Associations between pathway activation and African ancestry were calculated using the PROGENy model implemented in the *decoupleR* R package (version 2.12.0). The differential expression *t*-statistic calculated by the *Deseq2* R package was used as input in this analysis.

#### Cell Lineages Proportion Analysis

To investigate differences in cell lineage distribution across samples, we performed deconvolution analysis using the R package MuSiC^140^ (version 1.0.0), with a single-cell reference dataset derived from similar endometrial tissue. MuSiC was used to estimate the proportions of six lineage types: endothelial, epithelial, lymphocyte, epithelial/stromal, mesenchymal, and myeloid lineages. The epithelial/mesenchymal ratio was calculated by dividing the estimated epithelial proportion by the estimated mesenchymal proportion plus 1 to avoid division by zero. As MuSiC requires count data, we used the batch-corrected count matrix for this analysis.

Estimated lineage proportions were then compared across molecular subtypes. Samples with unclassified molecular subtypes (CBD) and samples with clear cell carcinoma histology were excluded from the analysis. CBD samples were excluded due to their high heterogeneity, and clear cell carcinoma samples were excluded due to the low sample size, which limited statistical power. For the remaining subtypes, molecular subtype annotation was retained as originally defined, except for the high copy number subtype (CN-H), which was further stratified into two groups based on histology with one group comprising endometrioid and serous cases while the other consisting of carcinosarcoma cases.

To assess associations between ancestry and estimated lineage proportions, we evaluated whether lineage proportions were associated with percent African (AFR) ancestry. Linear regression models were fitted with each lineage proportion as the outcome and either percent AFR ancestry as the predictor, adjusting for tumor purity and molecular subtype. Model coefficients represent the change in lineage proportion per one-unit increase in percent AFR ancestry.

## Data Availability

The genomic data are being deposited in the ICGC-ARGO data repository and will be made available upon publication.

## Code Availability

Custom analysis scripts will be made available in a GitHub repository upon publication

## Acknowledgements

The authors would like to thank the patients for their donation of tissues and clinical data. This study would not otherwise have been possible without their invaluable contribution. This work was conducted as part of New York Genome Center’s Polyethnic-1000 initiative. Funding and other external support for Polyethnic-1000 were provided in part by Illumina, Inc. Sample procurement, next-generation sequencing, and clinical data harmonization was performed by the New York Genome Center in collaboration with Northwell Health and Cold Spring Harbor Laboratory. This work was financially supported by grants to P.B and A.K from the National Institutes of Health (U01 CA289357) and to S.B from the National Cancer Institute (R37CA292807), Oliver S. and Jennie R. Donaldson Charitable Trust, the Mark Foundation for Cancer Research (20-028-EDV), Chan Zuckerberg Initiative/Silicon Valley Community Foundation (2021-239862), the Cold Spring Harbor Laboratory and Northwell Health Affiliation and Swim Across America. This work was performed with assistance from the US National Institutes of Health Grant S10OD028632-01. The results shown here are in part based upon data generated by the TCGA Research Network: https://www.cancer.gov/tcga.

## Author contributions

**Conceptualization**: MF, WRM, NR, SB, NC; **Data Curation:** MF, DG, ZG, WFH, KF, PB, MK, BY, TC, ZV, KO, ZS, LW, MB, NR, SB; **Formal Analysis:** DG, ZG, WFH, KF, AD, HG, PB, MK, TC, AO, ZV, VG, ALA, NR; **Investigation**: MF, DG, ZG, WFH, KF, AD, HG, PB, MK, BY, TC, AO, ZV, VG, CC, AN, OE, KO, MB; **Visualization:** DG, ZG, WFH, KF, AD, HG, MK, TC, AO, NR, NC; **Writing - Original Draft Preparation:** MF, DG, ZG, WFH, KF, AD, HG, MK, AO, MB, NR, NC; **Writing - Review and Editing**: MF, DG, WFH, KF, AD, MK, AO, AK, WRM, MB, NR, SB, NC; **Funding Acquisition**: MF, LW, NR, SB; **Resources** MF, SB; **Project Administration**: MF, ZS, LW, SB, NR, NC; **Supervision**: MF, WR,MC, SB, NR, NC. All authors reviewed and approved the final manuscript.

